# The distribution of global tidal marshes from earth observation data

**DOI:** 10.1101/2023.05.26.542433

**Authors:** Thomas A. Worthington, Mark Spalding, Emily Landis, Tania L. Maxwell, Alejandro Navarro, Lindsey S. Smart, Nicholas J. Murray

## Abstract

**Aim:** Tidal marsh ecosystems are heavily impacted by human activities, highlighting a pressing need to address gaps in our knowledge of their distribution. To better understand the global distribution and changes in tidal marsh extent, and identify opportunities for their conservation and restoration, it is critical to develop a spatial knowledge base of their global occurrence. Here, we develop a globally consistent tidal marsh distribution map for the year 2020 at 10-m resolution.

**Location:** Global

**Time period:** 2020

**Major taxa studied:** Tidal marshes

**Methods:** To map the location of the world’s tidal marshes we applied a random forest classification model to earth observation data from the year 2020. We trained the classification model with a reference dataset developed to support distribution mapping of coastal ecosystems, and predicted the spatial distribution of tidal marshes between 60°N to 60°S. We validated the tidal marsh map using standard accuracy assessment methods, with our final map having an overall accuracy score of 0.852.

**Results:** We estimate the global extent of tidal marshes in 2020 to be 52,880 km^2^ (95% CI: 32,030 to 59,780 km^2^) distributed across 120 countries and territories. Tidal marsh distribution is centred in temperate and Arctic regions, with nearly half of the global extent of tidal marshes occurring in the temperate Northern Atlantic (45%) region. At the national scale, over a third of the global extent (18,510 km^2^; CI: 11,200 – 20,900) occurs within the USA.

**Main conclusions:** Our analysis provides the most detailed spatial data on global tidal marsh distribution to date and shows that tidal marshes occur in more countries and across a greater proportion of the world’s coastline than previous mapping studies. Our map fills a major knowledge gap regarding the distribution of the world’s coastal ecosystems and provides the baseline needed for measuring changes in tidal marsh extent and estimating their value in terms of ecosystem services

## 1. INTRODUCTION

Tidal marsh ecosystems are vegetated coastal wetlands located in areas of regular to occasional tidal inundation, formed by a broad variety of herbaceous and woody vascular plants (Adam, 2002). Whilst dominated by saline and brackish marshes, they include areas of freshwater tidal marsh (Barendregt, 2018; Mitsch & Gosselink, 2015). They occur along many of the world’s sheltered, sediment-dominated coastlines, particularly in temperate and arctic regions (Tiner & Milton, 2018). They are also an important, yet frequently overlooked, coastal ecosystem in many arid and tropical regions (Viswanathan et al., 2020). Tidal marsh vegetation occurs between mean high water neaps up to the limits of the highest astronomical tide (Adam, 2002), often with distinct zonation across the tidal frame (Mitsch & Gosselink, 2015; Scott et al., 2014b, 2014a; Tiner & Milton, 2018). Tidal marshes show close ecological and physical linkages with adjacent coastal ecosystems, including unvegetated intertidal flats, shellfish beds and reefs, and seagrass beds lower in the tidal frame (Adam, 2002). They often form part of a more mosaicked habitat with mangroves or salt pans, particularly in warm temperate to tropical regions (Lopez-Portillo & Ezcurra, 1989; Rodriguez et al., 2016).

Tidal marshes are subject to a multitude of anthropogenic pressures, primarily because they are often located close to the most densely populated parts of the planet (Neumann et al., 2015). Their loss and degradation have been caused by a range of factors, tracing back over centuries and even millennia (Airoldi & Beck, 2007; Allen, 2000). These include land reclamation for conversion to agriculture and coastal infrastructure (Davy et al., 2009; Gedan & Silliman, 2009; Gu et al., 2018; Melville et al., 2016; Shi-lun & Ji-yu, 1995), aquaculture and salt production (Almeida et al., 2014), and invasion of *Spartina alterniflora* outside its native range (Zheng et al., 2018). In addition, tidal marshes are likely to be impacted by the multifaceted threats linked to climate change, including sea level rise and associated coastal squeeze, increased magnitude and frequency of extreme weather events and changes in precipitation and temperature (Adams, 2020; Silliman et al., 2005). The evidence of climate change impacts has already been detected with the reduction in extreme cold events allowing the poleward expansion of mangroves into tidal marsh habitat (Cavanaugh et al., 2014, 2019).

Recently, tidal marshes, alongside other coastal wetlands such as mangroves and seagrass, have garnered significant attention for their conservation value and restoration potential. In addition to their significance for biodiversity, tidal marshes have been recognised as supporting multiple ecosystem services, including carbon sequestration (Friess et al., 2020; zu Ermgassen et al., 2021). Tidal marshes are highly productive, with sequestration rates greater than some other terrestrial ecosystems (on average 210 g CO_2_ m^-2^ yr^-1^) (Chmura et al., 2003; Hopkinson et al., 2012), with potential global carbon stocks in tidal marsh soil estimated between 862 and 1,350 Tg C (Macreadie et al., 2021). Tidal marsh vegetation also helps attenuate wave energy (Möller et al., 2014) therefore providing storm protection benefits to coastal communities (Costanza et al., 2008; Shepard et al., 2011). They are important areas for tourism and recreation (Barbier et al., 2011) and they help support healthy fisheries that provide food and income for millions of people (Baker et al., 2020; Jänes et al., 2020). However, our ability to comprehensively estimate the value of tidal marshes and develop coordinated strategies for their protection and restoration at large scales is hindered by limited knowledge of their global distribution.

Despite the fact that tidal marshes can be identified from satellite images (Tiner & Milton, 2018), unlike other coastal ecosystems such as coral reefs (Li et al., 2020), mangroves (Bunting et al., 2022; Giri et al., 2011), and tidal flats (Murray et al., 2019), spatial mapping of tidal marshes has generally been conducted at local or regional scales. Until now there has been no globally consistent map of tidal marshes, and their total extent has been poorly quantified as a result (McLeod et al., 2011; Pendleton et al., 2012).

Efforts to map tidal marshes have utilized different methods, across different time periods, making comparisons across time and space unreliable. Where larger scale assessments exist, they tend to focus on well-studied regions such as the U.S.A. (U. S. Fish and Wildlife Service, 2021), China (Hu et al., 2021), Australia and parts of South America (Isacch et al., 2006). The most recent global assessment of tidal marsh spatial distribution collated Geographic Information System (GIS) data from peer-reviewed articles and grey literature, including spatial data from government agencies, non-governmental organisations and research institutions (Mcowen et al., 2017). In total, the authors identified almost 55,000 km^2^ of tidal marshes across 43 countries and territories, although they also identified several large regions – Canada, Northern Russia, South America and Africa – for which data was lacking (Mcowen et al., 2017). A global assessment of all wetlands, Zhang *et al*. (2023) also included a category of tidal salt marsh, mapping approximately 75,000km^2^, with other estimates from the literature ranging from 22,000 – 400,000 km^2^ (McLeod et al., 2011; Pendleton et al., 2012). Other efforts to characterise tidal marsh dynamics globally have typically focused on change analyses (Campbell et al., 2022; Murray, Worthington, et al., 2022) or simulation (Schuerch et al., 2018); however, a high-resolution map baseline has yet to be established.

Here, we create a globally consistent tidal marsh distribution map for the year 2020 at 10-m resolution. To do this we leverage the extensive archive of analysis-ready, publicly available remote-sensing imagery, fusing together optical and radar images from the European Space Agency’s Sentinel missions (Berger et al., 2012). This is coupled with an open-access global repository of global training and validation data (Murray, Bunting, et al., 2022). This wealth of data was analysed using the high-performance processing capabilities of the Google Earth Engine platform, which supports rapid planetary-scale remote-sensing data analyses (Gorelick et al., 2017). The tidal marsh distribution dataset provides a critical baseline for future analyses of changes in tidal marsh distribution and condition, and valuation of tidal marsh ecosystem services.

## 2. MATERIAL AND METHODS

To map the distribution of the world’s tidal marshes to allow estimates of extent and underpin future analyses of tidal marsh change, we developed a supervised classification of globally comprehensive earth observation data. We used 10 metre resolution active (Sentinel 1) and passive (Sentinel 2) data acquired in 2020 (Berger et al., 2012) and an open access reference dataset (Murray, Bunting, et al., 2022) to parameterise a random forest classification model that enabled the prediction of tidal marsh occurrence over the study area.

### 2.1 Covariate data

To develop a covariate set suitable to support the classification model, we filtered all images in the Sentinel 1 and 2 archives for the year 2020, retaining all images within our mapping area. The mapped area was confined between 60°N to 60°S, and within a coastal zone data mask (after Murray, Worthington, et al., 2022). In total, 287,320 optical and 143,067 radar images were processed.

The Sentinel 2 optical data was filtered to remove images with ≥ 20% cloudy pixels, and further masks were applied to remove pixels flagged in image metadata as clouds, cloud shadow or snow. Three spectral indices that represent either vegetation or water dynamics, and are therefore useful to discriminate tidal marshes from other land classes in a classification model, were calculated for every image. The normalized difference vegetation index (NDVI) is the normalized ratio of the near infrared (NIR) band which is reflected by vegetation and the red band which is absorbed by vegetation (Pettorelli et al., 2005).

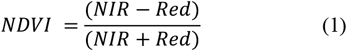

Values range between -1 and +1, with values closer to +1 representing areas of dense green vegetation. The enhanced vegetation index (EVI) was developed to reduce influence of atmospheric conditions and decouple the canopy background signal (Huete et al., 2002). In addition to the NIR and red bands used in NDVI, EVI uses the blue band to reduce the impact of atmospheric effects (Schultz et al., 2016).

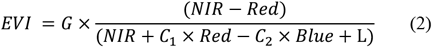

The coefficient L is the canopy background adjustment and C_1_ and C_2_ are used with the blue band to reduce aerosol influences on the red band (Huete et al., 2002). Values of L = 1, C_1_ = 6, C_2_ = 7.5, and G = 2.5 were used (Huete et al., 2002). The automated water extraction index (AWEI) combines the green, NIR and short-wave infrared (SWIR) bands to identify areas of water (Feyisa et al., 2014).

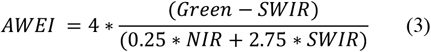

An annual composite of the optical images was created by taking the median, the 10^th^, 25^th^, 75^th^, and 90^th^ percentiles, the standard deviation, and the 5^th^ – 95^th^, 10^th^ – 90^th^, and 25^th^ – 75^th^ interval means of the NDVI, EVI and AWEI spectral indices and the raw NIR band, resulting in an initial set of 36 covariates. Exploratory analysis of 41,138 training points revealed high collinearity between the 36 covariates derived from the optical data. *A priori* we retained the median and standard deviation covariates, and used the *findCorrelation* function from the ‘caret’ R package (Kuhn, 2021) in R to remove highly correlated covariates from the remaining 28 covariates. Based on a threshold of r < 0.9, we removed 17 highly correlated covariates from the covariate set, resulting in a final set of 19 covariates for the classification model (Supporting Information Table S1). Annual composite metrics were used to reduce the impact of clouds, cloud shadow or snow on the indices, represent complex spectral dynamics in a manner suitable for a pixel-based classification model, and have been shown to be effective in estimating the distribution of a range of other coastal ecosystem types (Murray et al., 2019; Murray, Worthington, et al., 2022).

The Sentinel 1 radar data was filtered to those images with VV and VH polarizations, and indices that represented the span and total of the scattering power, the difference between co- and cross-polarized observations and the ratio of the VV and VH polarizations (Mahdianpari et al., 2019), were calculated for each image. An annual composite of the radar images was created by taking the median of the raw VV and VH polarizations and the Span, Difference and Ratio indices, resulting in five covariates (Supporting Information Table S1).

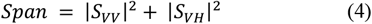

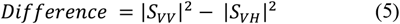

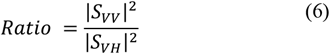

In addition to the radar and optical data, three covariates were included to assist in differentiating tidal marshes from other land classes that occur in coastal environments. Firstly, a global map of the probability of occurrence of tidal wetlands derived from over 1.1 million satellite images acquired by Landsat 5 to 8 (Murray, Worthington, et al., 2022) was used to inform the potential distribution of tidal marsh ecosystems. A global elevation model that combines land topography and ocean bathymetry (Amante & Eakins, 2009) was used to inform the classification model about real features of coastal environments that influence the distribution of tidal marshes, while an annual composite of monthly nightlights data from the year 2020 (Elvidge et al., 2017) was included to assist differentiation of urban and industrial areas.

The optical and radar data, and the three additional covariates (Supporting Information Table S1) were combined into a single composite image. Superpixel clustering based on the Simple Non-Iterative Clustering image segmentation algorithm was then applied to the composite image (Achanta & Süsstrunk, 2017). The image segmentation clusters areas of homogenous pixels and helps to reduce false positives in terms of individual pixels of the prediction class being predicted across the image. The pixel values for each band within a cluster are converted into the mean value of the band across the cluster, and as such the prediction of each pixel within a cluster results in the same classification. Such object-based approaches have been shown to have a superior predictive power to pixel-based classification when mapping coastal environments (Lyons et al., 2020; Mahdianpari et al., 2019).

### 2.2 Training data

To parametrise the random forest classification model, we collated a training dataset consisting of the known locations of five classes: tidal wetland ecosystems (tidal marshes, mangroves, and tidal flats) and non-tidal wetland types (permanent water and other terrestrial areas). The training set was assembled from three sources. Firstly, 136,404 points from the *coastTrain* dataset (Murray, Bunting, et al., 2022), which was created to provide reference data for coastal ecosystem remote sensing classification models. Secondly, a further 8,789 points developed to map coastal ecosystems in Australia (A. Navarro, personal communication). Finally, 754 tidal marsh locations collected specifically for this analysis. The 754 tidal marsh points were targeted at regions that had reduced coverage in the coastTrain dataset, such as South Africa, East Asia (outside China), the Middle East and Pacific coast of South

America. Potential locations of tidal marshes in these regions were identified from published literature and online sources. Visual interpretation of high-resolution satellite images available from Google Earth, alongside NIR and false colour composites from Landsat imagery were used to select pixels that represented tidal marshes. The combined training dataset consisted of a high number of mangrove locations from the coastTrain dataset, therefore a random sample of only 10,000 of the mangrove points was used to balance the training data. This resulted in a final training dataset of 41,762 points, including 9,811 tidal marsh locations (Supporting Information Figure S1).

### 2.3 Tidal marsh distribution model

To map the global distribution of tidal marshes we developed a random forest classification model. Machine learning approaches such as random forest have been applied to a variety of ecological and remote sensing topics due to their high classification accuracy and ability to rapidly model large datasets with complex interactions between predictor variables (Belgiu & Drăgu, 2016; Cutler et al., 2007). Machine learning approaches have been effectively applied to mapping coastal ecosystems such as intertidal wetlands (Murray, Worthington, et al., 2022), tidal flats (Murray et al., 2019), mangroves (Bunting et al., 2018) and coral reefs (Lyons et al., 2020).

#### 2.3.1 Model tuning

The training data was combined into two classes, tidal marshes and non-tidal marshes (permanent water, other terrestrial areas, mangroves, and tidal flats). Before parametrising the model in Google Earth Engine, we firstly tuned the random forest hyper-parameters using iterative testing of a hypergrid of potential values. Models were fitted using the training data and R package ‘ranger’ (Wright & Ziegler, 2017). The hypergrid consisted of 375 potential model parameterisations with differing values for the number of trees grown, the number of covariates sampled at each split, the fraction of observations sampled at each split, and the minimum node size. To account for minimal differences in the model fits (based on out-of-bag error rate) between the best fitting parametrizations, the median of the top ten models based on the hypergrid search was used in Google Earth Engine.

An initial model was fitted to the training data in Google Earth Engine with a random 80:20% training:validation split. Exploratory analysis suggested high overall accuracy (96.4%) on the validation dataset, with a Kappa coefficient of 0.90, and a producer’s accuracy for the different classes of 0.98 non-tidal marsh, 0.91 tidal marsh, and a user’s accuracy for the different classes of 0.97 non-tidal marsh, 0.94 tidal marsh. Owing to this high initial model accuracy, a final random forest classification model was then fitted in Google Earth Engine using all the training data. The final random forest classification model was then applied to the segmented composite image (containing the optical, radar and three additional covariates, Supporting Information Table S1), and each pixel was classified as either tidal marsh or not.

#### 2.3.2 Post-processing

The initial predicted distribution of tidal marshes was then refined using a post-processing procedure consisting of several steps. Areas that were predicted to have a low probability (<50%) of being a coastal wetland based on the work of Murray *et al*. (2022) were removed, as were areas greater than 10m in elevation using the MERIT Digital Elevation Model (Yamazaki et al., 2017) and those areas that overlapped with the predicted 2020 mangrove distribution (Bunting et al., 2022). The minimum mapping unit of the global tidal marsh map was tested for two areas (1-hectare versus 10-hectares) by identifying areas that had a minimum of either 100 or 1,000 connected pixels.

#### 2.3.3 Validation

This post-processed model prediction was validated using 2,300 randomly sampled points developed using the following stratified sampling procedure. Points were sampled equally across ten of the 11 biogeographical realms in the Marine Ecoregions of the World (Spalding et al., 2007). One hundred points were sampled from the non-tidal marsh class, as were a further 100 points classified as tidal marsh in both the 1-hectare and 10-hectare minimum mapping unit map versions. Finally, an additional 30 points classified as tidal marsh only in the 1-hectare minimum mapping unit version were sampled. A single image analyst used Google Satellite, Bing Maps, and Google Earth Pro imagery to assess each point of the validation set and assign it to one of three groups, ‘tidal marsh’, ‘other’, or ‘unknown’ using the Class Accuracy plugin in QGIS (Bunting, 2020). The model prediction for the point was concealed from the reviewer during the validation process. The ‘unknown’ assignment in the validation set was used for points where no confident assignment of the ecosystem type was possible, primarily due to poor reference imagery or a lack of information available about tidal marshes in particular regions, predominantly in parts of the tropics

Accuracy statistics were calculated for the different realms using the ‘caret’ R package (Kuhn, 2021) in R (R Core Team, 2021). Only validation points assigned to the groups ‘tidal marsh’ and ‘other’ were used to calculate the accuracy statistics (n = 1,708). For overall and producer’s accuracy, the validation statistics for the vast majority of realms were higher for the 10-hectare minimum mapping unit version (Supporting Information Table S2) in comparison to the 1-hectare minimum mapping unit version (Supporting Information Table S3), and this was therefore the version used for the final mapping product. Accuracy was very variable across the realms, with temperate regions generally achieving higher overall map accuracy. The user’s accuracy was always higher than the producer’s accuracy, which in the tropics was very low. To address the issues identified during the map validation, we undertook two final post-processing procedures. We removed areas of tidal marsh that had been identified as aquaculture ponds in ten countries in Asia, following Murray *et al*. (2022), and manually corrected misclassifications. Manual correction was carried out by visually assessing the map outputs and removing obvious misclassifications. Misclassifications were generally related to areas of aquaculture and flooded agriculture such as rice fields, and rocky shorelines.

The 1,708 previously classified validation points were then compared to the final version of the map, and accuracy statistics were again calculated. The manual corrections greatly improved the overall accuracy across the realms, with the largest improvements in tropical regions, although producer’s accuracy remained extremely in those regions (Table 1). We used resampling procedures to calculate the confidence intervals around our global accuracy statistics (Lyons et al., 2018), which were used to propagate uncertainty around derived estimates of global and regional tidal marsh extent. We resampled (n = 1,000 iterations) the validation points, using the mean of the samples as our estimates of accuracy, and the 0.025 and 0.975 percentiles to create the 95% confidence intervals (Supporting Information Table S4). Our final map had an overall accuracy of 0.852 (95% confidence interval (CI): 0.837 – 0.867). The resampling procedure allowed for asymmetry in the confidence intervals around our tidal marsh extent estimates, which better represent the unevenness in the omission and commission errors identified in our map (Supporting Information Tables S4-S5). We used the 0.05 percentile of the user’s and producer’s error estimates from the resampled distribution to calculate the upper and lower bounds of all area estimates using the formulas:

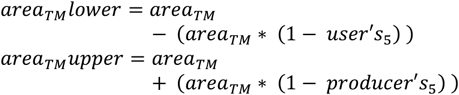

Where *area _™_* is the area of tidal marsh, *user’s*_5_ and *Producer’s*_5_ are the 0.05 percentile of the user’s producer’s error estimates, respectively.

**Table 1:** Realm level accuracy statistics for the final tidal marsh map. Producer’s and user’s accuracy statistics given for tidal marsh class only. The number of sample points (N) within each realm is the maximum number assessed (n = 200) minus those classified as unknown during the validation assessment.

| Realm | Overall Accuracy | Kappa | Producer's Accuracy | User's Accuracy | N |
| --- | --- | --- | --- | --- | --- |
| Temperate Northern Atlantic | 0.88 | 0.76 | 0.80 | 0.95 | 200 |
| Temperate Northern Pacific | 0.81 | 0.60 | 0.64 | 0.88 | 171 |
| Tropical Atlantic | 0.80 | 0.38 | 0.34 | 0.80 | 129 |
| Western Indo-Pacific | 0.73 | 0.13 | 0.11 | 0.71 | 151 |
| Central Indo-Pacific | 0.91 | 0.69 | 0.73 | 0.77 | 171 |
| Eastern Indo-Pacific | 0.93 | 0.43 | 0.32 | 0.86 | 190 |
| Tropical Eastern Pacific | 0.85 | 0.09 | 0.06 | 1.00 | 114 |
| Temperate South America | 0.80 | 0.59 | 0.63 | 0.90 | 188 |
| Temperate Southern Africa | 0.89 | 0.78 | 0.84 | 0.89 | 197 |
| Temperate Australasia | 0.88 | 0.75 | 0.82 | 0.92 | 197 |

#### 2.3.4 Extent statistics

Tidal marsh area was summarised at the marine ecoregion realm level (Spalding et al., 2007) and for countries and territories using the union of the ESRI world country database and the Exclusive Economic Zones version 11 (Flanders Marine Institute, 2020). The percentage of the tidal marsh contained within protected areas in each country was calculated using The World Database on Protected Areas (WDPA; UNEP-WCMC and IUCN, 2023). The WDPA was cleaned following standard procedures (Hanson, 2022), which included removing polygon vertices, removing protected areas where the status was proposed or unknown, and dissolving protected area polygons to prevent double counting of overlapping protected areas.

## 3. RESULTS AND DISCUSSION

Knowledge of the distribution of tidal marsh ecosystems is essential to understand fundamental drivers of their dynamics, estimate risks to their persistence, establish baselines for assessing a range of national-to-global conservation targets and to provide a basis for accounting for their ecosystem service provisioning. Our study reports the development of the first global, high-resolution thematic map of tidal marsh distribution, and allows a range of analyses that can fill these important knowledge gaps.

### 3.1 Global and regional estimates

We estimate the global extent of tidal marshes in 2020 to be 52,880 km^2^ (CI: 32,020 to 59,780 km^2^). This estimate is lower than many earlier estimates generated by diverse mapping approaches, and confirms that tidal marshes occupy a considerably smaller area than other coastal ecosystems such as, for example, mangroves 147,359 km^2^ (Bunting et al., 2022) and tidal flats 127,921 km^2^ (Murray et al., 2019).

Table 2 shows the estimates of tidal marsh extent by biogeographic realm (Spalding et al., 2007). The findings highlight the predominance of these ecosystems in the temperate and Arctic realms, and the particular importance of the temperate Northern Atlantic. This single realm hosts 45% of the world’s tidal marshes, with extensive areas along the Atlantic and Gulf of Mexico coasts of the U.S.A and in the Northern European Seas province (Figure 1). The widespread distribution of marshes in this region is likely to be a product of multiple geological and geomorphological factors influencing the abundance of extensive, protected and low-elevation coastal sediments. Climate too is important – it is too cold for mangroves, but not influenced by ice-scour. (Scott et al., 2014a). As noted by previous authors, this realm is also the centre of floristic diversity for tidal marshes (Adam, 1990; Chapman, 1974). We estimate that the temperate Northern Pacific Realm also has significant areas of tidal marshes, especially around the coasts of southern Alaska and the Russian coasts of the Sea of Okhotsk, an area that was unmapped by Mcowen *et al*. (2017).

**Table 2:** The extent of tidal marshes by biogeographic realm (Spalding et al., 2007). 95% confidence intervals created by resampling (n = 1,000) the validation points and using the 0.05 percentile of the user’s and producer’s error estimates.

| Realm | Area (km<sup>2</sup>) | 95% Confidence Interval | % of Global Total |
| --- | --- | --- | --- |
| Arctic* | 9210 | 5580 - 10410 | 17.4 |
| Central Indo-Pacific | 600 | 360 - 670 | 1.1 |
| Eastern Indo-Pacific | <10 | - | <0.01 |
| Southern Ocean | - | - | - |
| Temperate Australasia | 2040 | 1230 - 2300 | 3.9 |
| Temperate Northern Atlantic | 23760 | 14390 - 26860 | 44.9 |
| Temperate Northern Pacific | 6580 | 3980 - 7440 | 12.4 |
| Temperate South America | 3790 | 2300 - 4290 | 7.2 |
| Temperate Southern Africa | 80 | 50 - 100 | 0.2 |
| Tropical Atlantic | 4810 | 2920 - 5440 | 9.1 |
| Tropical Eastern Pacific | 80 | 50 - 90 | 0.2 |
| Western Indo-Pacific | 1920 | 1170 - 2180 | 3.6 |
| **Total** | **52880** | **32020 - 59780** |  |

\*N.B. our map does not extend beyond 60°N and 60°S therefore the total area for the Arctic realm is an underestimate. For the Southern Ocean there are no records of tidal marshes on mainland Antarctica (Greenberg et al., 2006), and our map does not classify any of the limited coastal wetlands on the sub-Antarctic islands as tidal marshes.

**Figure 1:**
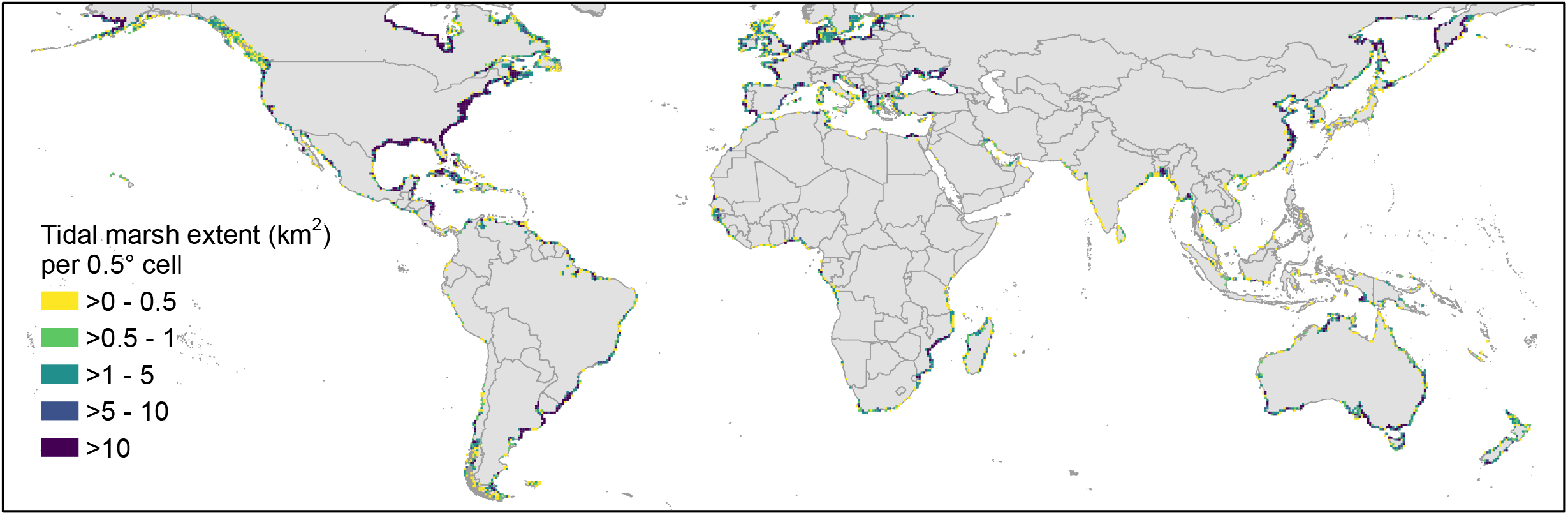
The 2020 distribution of tidal marshes, with darker colours representing greater extents of tidal marshes (km^2^) within a 0.5° grid cell.

The area of tidal marshes in the southern hemisphere is 8,380 km^2^ (CI: 5,080 to 9,480 km^2^) representing only 16% of the full extent of coastal tidal marsh we detected in our analysis. However, the Atlantic coast of South America in particular supports extensive tidal marshes on estuaries with large discharges (Hatje et al., 2023): Isacch et al. (2006) estimated some 2,133 km^2^ of tidal marshes in the area between southern Brazil and central Argentina, which is lower than our estimate of 3,060 km^2^ for that region. Our prediction for Temperate Southern Africa confirms prior observations that these are indeed scarce habitats in this realm (Adams, 2020).

Our map indicated tropical realms support 7,410 km^2^ (CI: 4,500 to 8,380 km^2^) of tidal marshes. In the tropics, coastal wetlands are typically dominated by mangroves, however tidal marshes are still found in most areas (Almahasheer, 2021; e.g., Hena et al., 2007; Viswanathan et al., 2020), albeit often within spatially confined areas. Viswanathan *et al*. (2020) estimated 290 km^2^ of ‘salt marsh’ around the coast of India, considerably more than the 50 km^2^ from our analysis, however, the density of salt marsh vegetation in the Indian study (19 ± 1 plants per m^2^) was lower than the average density of tidal marshes in temperate regions, highlighting the challenge of identifying tidal marshes with different structures in different regions of the world. Within the Indo-Pacific Realms our maps show larger distribution of tidal marsh around the coasts of Mozambique, Madagascar and northern Australia. These are somewhat arid macrotidal regions where tidal marshes typically occur behind mangroves, high in the tidal frame (Saintilan, 2009). In the Tropical Atlantic, tidal marshes are widespread – notably in Central America and Cuba – again in close proximity to mangroves, and usually in the upper reaches of the tidal frame.

### 3.2 National estimates

Political contexts set the scene for conservation action and hence understanding distribution statistics at national levels is important and serves to provide data for National Biodiversity Strategies and Action Plans and the United Nations’ System of Environmental Economic Accounting. Our map identifies tidal marsh ecosystems in 120 countries, and Figure 2 shows the extents for the 20 countries with the largest areas of tidal marsh. At the national scale, over a third of the global extent (18,510 km^2^; CI: 11,210 – 20,930) is estimated to be found within the U.S.A. (Figure 2), with tidal marshes across all coastlines, but most extensive on the Atlantic and Gulf of Mexico coasts. Canada (8,530 km^2^; CI: 5,170 – 9,650) and Russia (5,140 km^2^; CI: 3,110 – 5,810) are also important, and the combined extent of just these three countries makes up over 60% of global tidal marsh extent. Any future inclusion of data from above 60°N would further highlight this dominance. An additional six countries are estimated to have tidal marsh extents in excess of 1,000km^2^ (Supporting Information Table S6).

**Figure 2:**
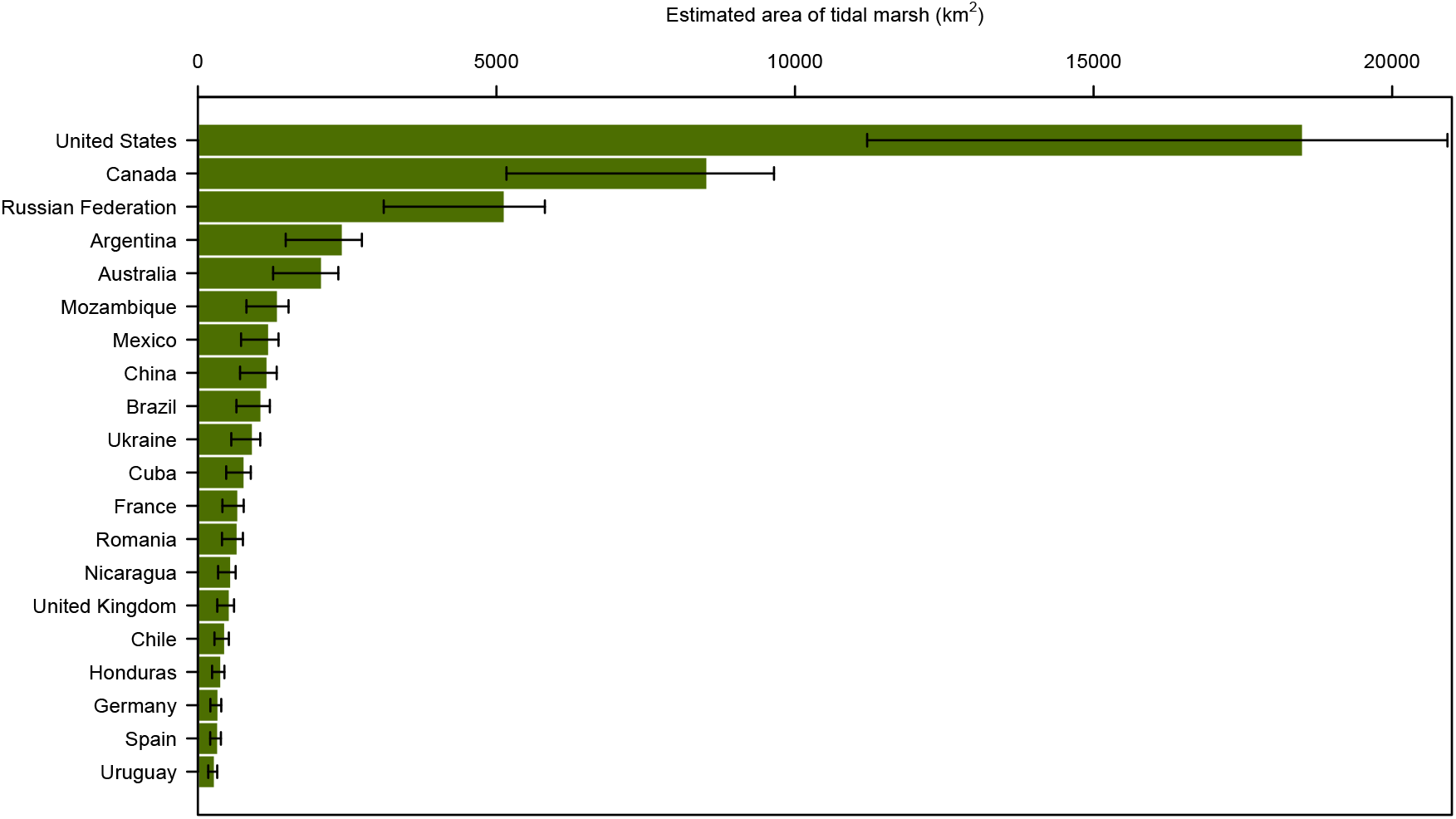
The area of tidal marsh in the countries with the largest extents. 95% confidence intervals were estimated by resampling the validation set (n = 1,000) and using the 0.05 percentile of the user’s and producer’s error estimates. The resampling procedure allowed for asymmetry in the confidence intervals around our tidal marsh extent estimates, which better represent the unevenness in the omission and commission errors identified in our map.

In Europe, tidal marshes are extensive, with a combined total of ∼5,000 km^2^, concentrated along the Atlantic coasts, the North and Baltic Seas. With its microtidal regime, and influenced by millennia of human conversion (Airoldi & Beck, 2007), the Mediterranean has only limited tidal marsh areas and many are fragmented. Our maps also show tidal marsh in the northern Black Sea, particularly in the major river deltas of the Danube and Dnepr and along the eastern shore of the Azov Sea (Figure 1).

### 3.3. Comparison to other global estimates

While our total extent closely matches the global estimate of Mcowen *et al*. (2017; 54,951 km^2^), there are notable differences in the distribution between the two maps. We identify tidal marshes in an extra 17 countries, while our maps include spatial extents for a further 71 countries where the Mcowen *et al*. (2017) provides only point locations. Our extent is also in line with a 45,000 km^2^ estimate for non-arctic tidal marshes (Greenberg et al., 2006) However, our extent is 58% - 71% of the estimates of Murray *et al*. (90,800km^2^; 2022) and Zhang *et al*. (74,910km^2^; 2023), respectively. The former was a more broadly based study of intertidal wetland dynamics and thus not directly comparable. Zhang *et al*. (2023) is a lower resolution (30m) global assessment of all wetlands, which includes “coastal saltmarsh”. The larger extent predicted is somewhat explained by the inclusion of 4,800 km^2^ along Arctic coastlines which were precluded from our study (see below). In addition, Zhang *et al*. (2023) study predicts the presence of “coastal saltmarsh” over a far greater proportion of the global coastline, however this is partly explained by the fact that it does not apply an area filter to remove the noise (predictions of wetland distribution confined to single pixels) in their maps. No recent maps have come close to the highest previous global estimates of 400,000 km^2^ (Duarte et al., 2005).

The strength of our approach is that it was targeted to the mapping of tidal marshes using a training sets that conform to a single class definition (e.g., Keith et al., 2022) and was developed via deep literature review and image interpretation. The image covariates were designed for the purpose of mapping tidal marshes (not all wetlands), and underpin a classification approach shown to perform well in coastal environments which is essential for resolving the uncertainty in tidal marsh distribution.

### 3.4 Protected Areas

We estimate that over 45% (24,200 km^2^) of the world’s 52,880 km^2^ of tidal marshes are found within the boundaries of protected areas. This is approximately equivalent to the extent of mangroves that occur within protected areas (42%; Leal & Spalding, 2022) and greater than the current area of coral reefs (32%; Marine Conservation Institute, 2018) or tidal flats (31%; Hill et al., 2021). Such extensive protection of coastal ecosystems likely reflects a growing perception of their immense value for biodiversity and for people. In this regard, tidal marshes are among the few ecosystem types globally that already meet the targets established by the Kunming-Montreal Global Biodiversity Framework with its commitment to conserve 30% of land and sea through the establishment of protected areas by 2030 (CBD, 2022), although it is important to be aware that if we could measure original extent, this figure would be considerably lower. The USA and Canada are critical in securing these levels of coverage with some 6,600 km^2^ and 3,500 km^2^ in protected areas, or 35% and 41% of their national tidal marsh extent respectively (Figure 3). In Europe the proportional coverage is particularly high, with 14 European countries having >90% of their extant tidal marshes within their protected area network (Figure 3; Supporting Information Table S6), although it is noteworthy that our knowledge of losses in this region suggest that most countries have lost between 50-90% of historical cover (Airoldi & Beck, 2007).

**Figure 3:**
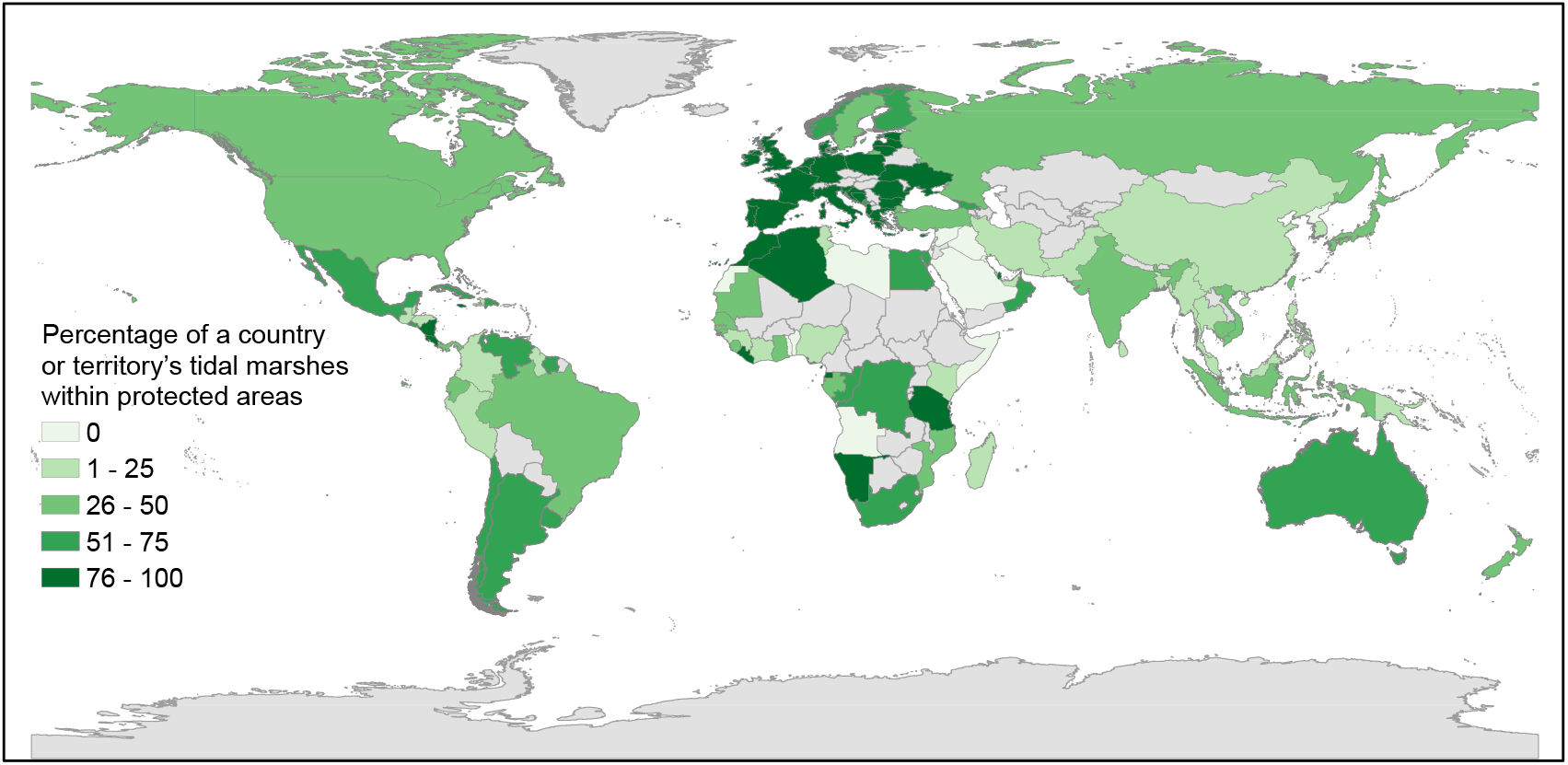
The percentage of the country or territory’s tidal marsh extent within the boundaries of a protected area.

### 3.5 Challenges and Limitations

One of the main challenges in developing a global map of tidal marshes is the considerable variability in an ecosystem that is globally distributed, ranging from monospecific reedbeds, salt-tolerant grasses and low, succulent shrubs, and that exists under highly variable environmental settings (Keith et al., 2022). The habitats captured in our analysis maps are highly diverse: many are extensive near-continuous areas dominating the upper reaches of wide intertidal frames. Elsewhere they are part of a tight mosaic with other habitats such as mangroves in tropical regions; or in complexes of dunes, lakes, marshes and drylands in the higher latitudes. In most areas tidal flushing is likely to be regular, but in some settings, such as in microtidal regimes, and in intermittently closed and open lagoons, such inundation may be more seasonal or intermittent.

In large part, the tidal marshes in our maps conform to the definition of saltmarsh developed by Adam (2002), which is close to that of Keith *et al*. (2022) for “Coastal saltmarshes and reedbeds”. By contrast, the Ramsar Convention recognizes several habitat types within tidal marshes including salt marshes, salt meadows, saltings, tidal brackish and freshwater marshes (Department of Climate Change Energy the Environment and Water, 2021). Even so, any mapping practise of this sort is in part constrained by what can be distinguished remotely from imagery or associated interpretation. In this study, our mapped distribution includes a considerable number of brackish to freshwater tidal marshes alongside the more strongly halophytic communities that have been described by others. This is a potential limitation of the research as it is currently not possible to differentiate such systems based on salinity using remote sensing methods; however, it ensures that we incorporate the entirety of an ecotone that is rarely, if ever, a sharp boundary (Maltby & Barker, 2009; Phelan et al., 2011).

The use of a global training dataset (Murray, Bunting, et al., 2022) was intended to include tidal marsh ecosystems present across all latitudes and representing very different tidal and climatic regimes; however, training data for the model was most comprehensive in areas where more tidal marshes occur, primarily temperate regions. This resulted in a potential bias in our prediction towards tidal marshes with physical features reflected by the training set and, aside from showing the distributional range of tidal marshes, it is noteworthy that our map also shows gaps – stretches of coastlines or regions where tidal marshes are rare to absent. These include many oceanic islands, but also some wet tropical regions in Southeast Asia. Gaps associated with hypersaline flats of northern Australia and the Middle East are also notable – these areas may have sparse saltmarsh plants fringing the margins of sabkha ecosystems (Barth & Böer, 2002), but these areas of very sparse fringing marsh vegetation were specifically not included in the training set.

The latitudinal limits to our study do not affect the southern hemisphere where tidal marsh is thought not to occur on the Antarctic continent (Greenberg et al., 2006); however, they mean we underestimate the overall tidal marsh extent in the Arctic. Our map only covers the southern parts of this climate region in Canada, USA, and Russia; the areas that likely to host a large proportion of arctic tidal marshes because they are less impacted by extreme conditions of temperature, snow and ice cover, ice scour, high wave energy and isostatic uplift compared to the higher latitude coasts. Despite the gap in our coverage, the very large areas we show, notably in southern Baffin Bay in Canada, mean that this realm has the second highest tidal marsh coverage, globally. Moving north, beyond the reach of our maps, it is likely that tidal marshes will be less extensive owing to the impact of ice abrasion and the harsh climate (Adam, 2002; Scott et al., 2014a). Zhang *et al*. (2023) map 4,800 km^2^ in the high Arctic area, however given the differences in our mapping approaches, we would predict a much smaller extent than this in these high latitudes. It is important to note that while the tidal marshes in these high northern latitudes tend to be limited in distribution with low diversity, they are extremely vulnerable to environmental change (Adam, 1990, 2002; Chapman, 1974; Martini et al., 2019; Sergienko, 2013).

## 4. CONCLUSIONS

This research presents the first consistent high resolution tidal marsh distribution map for the world. Compared to previous studies, it identifies tidal marshes in more countries and across a greater proportion of the world’s coastline, and provides the highest-resolution view of the extant distribution of this globally widespread coastal ecosystem. While there are opportunities to improve the map, particularly in the high Arctic and some tropical and arid regions, it already provides an invaluable baseline against which to measure change and to quantify the value of important ecosystem services.

Historic losses of tidal marshes have been considerable, with some areas continuing to experience land use conversion and tidal marsh degradation. With a globally consistent method that focuses solely on tidal marshes to develop a strong baseline, it should be possible to use our map to identify recent losses, following the methods established for wetland ecosystems (Bunting et al., 2022; Campbell et al., 2022; Murray, Worthington, et al., 2022). Our map will therefore enable better tracking of ongoing changes, which may represent natural dynamics, the impacts of sea level rise (Saintilan et al., 2022), or direct human modifications including tidal marsh conversion or the impact of marsh restoration efforts (Murray, Worthington, et al., 2022). Other work is ongoing to model the carbon stocks in tidal marsh soils and coastal protection functions, while it is hoped that others may be able to develop improved maps of fisheries enhancement and other benefits. Tidal marshes, although spatially limited, represent ecosystems of critical importance to biodiversity and to people. By establishing an open-access, global baseline, we hope to encourage greater efforts to secure a long-term future for these ecosystems, and indeed for the millions of people who depend upon them.

## ACKNOWLEDGEMENTS

This project benefited from funding from the Bezos Earth Fund and other donors supporting the Nature Conservancy. N.J.M. was supported by an Australian Research Council Discovery Early Career Research Award DE190100101. For the purpose of open access, the author has applied a Creative Commons Attribution (CC BY) licence to any Author Accepted Manuscript version arising from this submission.

## DATA ACCESSIBILITY STATEMENT

The tidal marsh distribution data are viewable at https://tomworthington81.users.earthengine.app/view/global-tidal-marsh-distribution and made available directly via Google Earth Engine (Asset ID: users/tomworthington81/SM_Global_2020/global_export_v2_6/saltmarsh_v2_6).

