## Supporting Information for "The distribution of global tidal marshes from earth observation data"

**Table S1: Datasets used in the tidal marsh remote sensing classification model.**

| **Input Data** | **Variables** | **Reducers Applied** | **Number of Covariate Layers** |
|---|---|---|---|
| Sentinel-2 MSI: MultiSpectral Instrument, Level-2A | NDVI | Median<br>Standard deviation<br>$10^{th}$ percentile<br>$90^{th}$ percentile | 4 |
| | EVI | Median<br>Standard deviation<br>$10^{th}$ percentile<br>$25^{th}$ percentile<br>$90^{th}$ percentile<br>$10^{th}$ – $90^{th}$ interval means<br>$25^{th}$ – $75^{th}$ interval means | 7 |
| | AWEI | Median<br>Standard deviation<br>$25^{th}$ percentile<br>$75^{th}$ percentile | 4 |
| | NIR | Median<br>Standard deviation<br>$10^{th}$ percentile<br>$75^{th}$ percentile | 4 |
| Sentinel-1 SAR GRD: C-band Synthetic Aperture Radar | VV<br>VH<br>Span<br>Difference<br>Ration | Median | 5 |
| Murray Global Tidal Wetland Change v1.0 | Probability of occurrence of tidal wetlands | N/A | 1 |
| ETOPO1: Global 1 Arc-Minute Elevation | Elevation | N/A | 1 |
| VIIRS Nighttime Day/Night Band Composites Version 1 | Average DNB radiance values. | N/A | 1 |

**Table S2: Realm level accuracy statistics for the 10-hectare minimum mapping unit version of the tidal marsh model. Producer's and user's accuracy statistics given for tidal marsh class only. The number of sample points (N) within each realm is the maximum number assessed (n = 200) minus those classified as unknown during the validation assessment.**

| **Realm** | **Overall Accuracy** | **Kappa** | **Producer's Accuracy** | **User's Accuracy** | **N** |
|---|---|---|---|---|---|
| Temperate Northern Atlantic | 0.88 | 0.76 | 0.80 | 0.95 | 200 |
| Temperate Northern Pacific | 0.81 | 0.60 | 0.64 | 0.90 | 171 |
| Tropical Atlantic | 0.76 | 0.32 | 0.30 | 0.80 | 129 |
| Western Indo-Pacific | 0.60 | 0.06 | 0.08 | 0.71 | 151 |
| Central Indo-Pacific | 0.65 | 0.25 | 0.31 | 0.77 | 171 |
| Eastern Indo-Pacific | 0.55 | 0.06 | 0.07 | 0.86 | 190 |
| Tropical Eastern Pacific | 0.80 | 0.06 | 0.04 | 1.00 | 114 |
| Temperate South America | 0.78 | 0.54 | 0.60 | 0.90 | 188 |
| Temperate Southern Africa | 0.80 | 0.60 | 0.68 | 0.89 | 197 |
| Temperate Australasia | 0.85 | 0.71 | 0.78 | 0.92 | 197 |

**Table S3: Realm level accuracy statistics for the 1-hectare minimum mapping unit version of the tidal marsh model. Producer's and user's accuracy statistics given for tidal marsh class only. The number of sample points (N) within each realm is the maximum number assessed (n = 230) minus those classified as unknown during the validation assessment.**

| **Realm** | **Overall Accuracy** | **Kappa** | **Producer's Accuracy** | **User's Accuracy** | **N** |
|---|---|---|---|---|---|
| Temperate Northern Atlantic | 0.80 | 0.60 | 0.67 | 0.96 | 230 |
| Temperate Northern Pacific | 0.75 | 0.51 | 0.56 | 0.92 | 194 |
| Tropical Atlantic | 0.65 | 0.18 | 0.20 | 0.80 | 150 |
| Western Indo-Pacific | 0.55 | 0.07 | 0.09 | 0.78 | 169 |
| Central Indo-Pacific | 0.59 | 0.19 | 0.27 | 0.79 | 193 |
| Eastern Indo-Pacific | 0.48 | 0.04 | 0.06 | 0.88 | 220 |
| Tropical Eastern Pacific | 0.66 | 0.03 | 0.02 | 1.00 | 137 |
| Temperate South America | 0.72 | 0.46 | 0.54 | 0.91 | 217 |
| Temperate Southern Africa | 0.75 | 0.51 | 0.61 | 0.91 | 227 |
| Temperate Australasia | 0.79 | 0.59 | 0.69 | 0.93 | 227 |

**Table S4: Accuracy assessment statistics for the final version of the global tidal marsh map, based on a validation sample of 1,708 points. We resampled (n = 1,000 iterations) the validation points, using the mean of the samples as our estimates of accuracy and the 0.025 and 0.975 percentiles to create the 95% confidence intervals.**

| | | **95% Confidence Interval** | |
|---|---|---|---|
| **Metric** | **Estimate** | **Lower** | **Upper** |
| **Overall accuracy** | 0.852 | 0.837 | 0.867 |
| **Other (Producer's)** | 0.961 | 0.950 | 0.972 |
| **Other (User's)** | 0.840 | 0.824 | 0.854 |
| **Tidal Marsh (Producer's)** | 0.641 | 0.599 | 0.680 |
| **Tidal Marsh (User's)** | 0.893 | 0.866 | 0.922 |

**Table S5: Class accuracy results for the global tidal marsh map based on 1,708 validation sample points.**

| | | Reference | |
|---|---|---|---|
| | | **Other** | **Tidal Marsh** |
| **Mapped** | **Other** | 1086 | 44 |
| | **Tidal Marsh** | 208 | 370 |

**Table S6: The extent of tidal marsh by country or territory (Flanders Marine Institute, 2020) rounded to the nearest 10km$^2$. 95% confidence intervals created by resampling (n = 1,000) the validation points and using the 0.05 percentile of the user's and producer's error estimates. The area and percentage of the country or territory's tidal marsh extent within the boundaries of a protected area. Only country or territory's with > 10 km$^2$ of tidal marsh reported.**

| Country or Territory | Area (km$^2$) | 95% Confidence Interval | Extent within Protected Areas (%) |
|---|---|---|---|
| United States | 18,510 | 11,210 – 20,930 | 6,550 (35) |
| Canada | 8,530 | 5,170 – 9,650 | 3,520 (41) |
| Russian Federation | 5,140 | 3,110 – 5,810 | 1,670 (32) |
| Argentina | 2,430 | 1,470 – 2,750 | 1,410 (58) |
| Australia | 2,080 | 1,260 – 2,350 | 1,140 (55) |
| Mozambique | 1,340 | 810 – 1,520 | 680 (50) |
| Mexico | 1,190 | 720 – 1,350 | 880 (73) |
| China | 1,170 | 710 – 1,320 | 270 (23) |
| Brazil | 1,070 | 650 – 1,210 | 480 (45) |
| Ukraine | 920 | 560 – 1,040 | 780 (85) |
| Cuba | 780 | 470 - 890 | 540 (69) |
| France | 680 | 410 - 770 | 650 (96) |
| Romania | 670 | 400 - 750 | 670 (100) |
| Nicaragua | 560 | 340 - 630 | 480 (86) |
| United Kingdom | 540 | 320 - 610 | 470 (88) |
| Chile | 460 | 280 - 520 | 230 (51) |
| Honduras | 390 | 240 - 440 | 30 (8) |
| Germany | 350 | 210 - 390 | 330 (97) |
| Spain | 340 | 210 - 390 | 320 (94) |
| Uruguay | 290 | 170 - 320 | 190 (65) |
| Estonia | 270 | 160 - 300 | 220 (83) |
| Japan | 270 | 160 - 300 | 110 (43) |
| Denmark | 260 | 160 - 300 | 260 (97) |
| Turkey | 250 | 150 - 290 | 110 (42) |
| New Zealand | 240 | 150 - 280 | 100 (40) |
| Italy | 230 | 140 - 260 | 200 (89) |
| Greece | 210 | 130 - 240 | 210 (96) |
| Bangladesh | 200 | 120 - 230 | 10 (5) |
| Netherlands | 180 | 110 - 210 | 170 (94) |
| Korea, Republic of | 180 | 110 - 200 | 20 (11) |
| Egypt | 180 | 110 - 200 | 110 (62) |
| Indonesia | 170 | 100 - 190 | 50 (29) |
| Venezuela, Bolivarian Republic of | 170 | 100 - 190 | 110 (66) |
| Latvia | 160 | 100 - 180 | 130 (80) |

| | | | |
|---|---|---|---|
| Poland | 160 | 100 - 180 | 150 (91) |
| Portugal | 160 | 100 - 180 | 150 (92) |
| Madagascar | 150 | 90 - 170 | 20 (14) |
| Ireland | 130 | 80 - 150 | 110 (81) |
| Myanmar | 120 | 70 - 140 | 20 (13) |
| Sweden | 120 | 70 - 130 | 50 (40) |
| Bahamas | 120 | 70 - 130 | 40 (36) |
| South Africa | 90 | 50 - 100 | 50 (53) |
| Korea, Democratic People's Republic of | 80 | 50 - 90 | <10 (0) |
| Albania | 80 | 50 - 90 | 70 (84) |
| Guinea-Bissau | 70 | 40 - 80 | 20 (26) |
| Georgia | 70 | 40 - 70 | 40 (57) |
| Belize | 60 | 40 - 70 | 10 (18) |
| Benin | 50 | 30 - 60 | 0 (0) |
| Tunisia | 50 | 30 - 60 | 10 (23) |
| Papua New Guinea | 50 | 30 - 60 | <10 (5) |
| Senegal | 50 | 30 - 60 | 20 (46) |
| India | 50 | 30 - 60 | 20 (33) |
| Croatia | 50 | 30 - 60 | 50 (94) |
| Colombia | 50 | 30 - 50 | 10 (15) |
| Morocco | 40 | 30 - 50 | 40 (86) |
| Suriname | 40 | 30 - 50 | 20 (57) |
| Gambia | 40 | 20 - 50 | 10 (13) |
| Haiti | 40 | 20 - 40 | <10 (1) |
| Lithuania | 40 | 20 - 40 | 30 (81) |
| Gabon | 30 | 20 - 40 | 10 (32) |
| Guyana | 30 | 20 - 40 | 10 (25) |
| Mauritania | 30 | 20 - 30 | 10 (37) |
| Algeria | 30 | 20 - 30 | 20 (93) |
| Montenegro | 30 | 20 - 30 | 20 (72) |
| Nigeria | 20 | 10 - 20 | <10 (3) |
| Sierra Leone | 20 | 10 - 20 | 10 (34) |
| Jamaica | 20 | 10 - 20 | 20 (88) |
| Bulgaria | 20 | 10 - 20 | 20 (97) |
| Thailand | 20 | 10 - 20 | <10 (2) |
| Dominican Republic | 20 | 10 - 20 | 10 (52) |
| Angola | 20 | 10 - 20 | 0 (0) |
| Ghana | 20 | 10 - 20 | 10 (43) |
| Cambodia | 20 | 10 - 20 | 10 (48) |
| Costa Rica | 20 | 10 - 20 | 10 (94) |
| Viet Nam | 10 | 10 - 10 | <10 (38) |
| Guatemala | 10 | 10 - 10 | <10 (19) |
| Iran, Islamic Republic of | 10 | 10 - 10 | <10 (14) |
| Peru | 10 | 10 - 10 | <10 (1) |

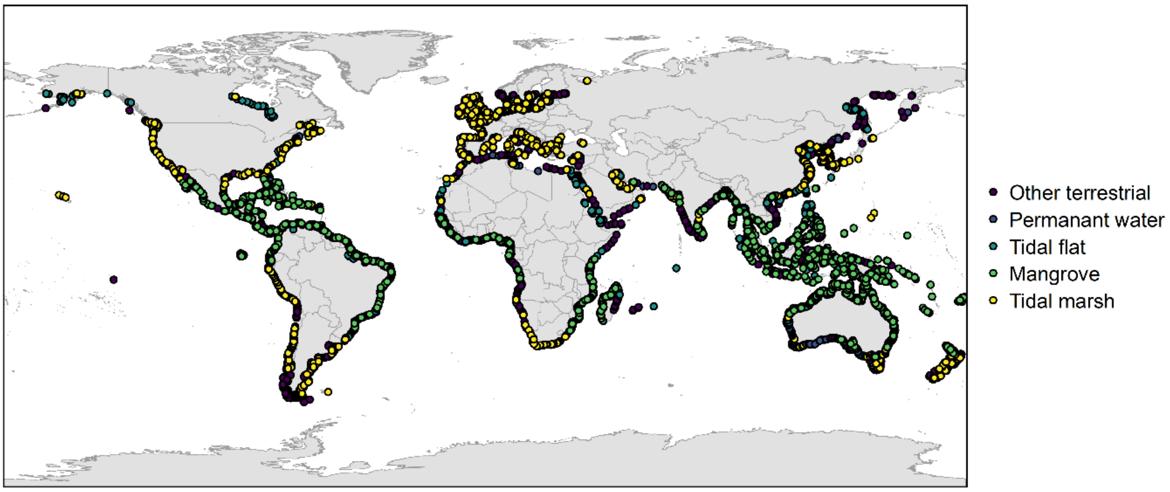


Figure S1: The distribution of the 41,762 points training points.
